# Oleic acid protects *Caenorhabditis* mothers from mating-induced death

**DOI:** 10.1101/2020.08.31.275198

**Authors:** Leo S. Choi, Cheng Shi, Coleen T. Murphy

## Abstract

Reproduction comes at a cost, including death. Previous studies of the interconnections between reproduction, lifespan, and fat metabolism in *C. elegans* were predominantly performed in low-reproduction conditions. To understand how increased reproduction affects lifespan and fat metabolism, we examined mated worms; we find that a Δ9 desaturase, FAT-7, is significantly up-regulated. Dietary supplementation of oleic acid (OA), the immediate downstream product of FAT-7 activity, restores fat storage and completely rescues mating-induced death, while other fatty acids cannot. OA-mediated lifespan restoration is also observed in *C. elegans* mutants suffering increased death from short-term mating, and in mated *C. remanei* females, indicating a conserved role of oleic acid in post-mating lifespan regulation. Because OA supplementation does not further increase the reproductive span or the brood size of mated *C. elegans* mothers, our results suggest that altering specific fat metabolism uncouples reproduction and somatic lifespan regulation, providing potent targets to ameliorate the cost of reproduction.

## Introduction

Reproduction is a costly process that depletes energy reserves that may be needed for somatic maintenance and survival (Williams, 1966; Hansen et al., 2013), thus linking reproduction with fat metabolism and lifespan regulation. In many organisms, increased reproduction leads to decreased lifespan, and vice versa (Drori and Folman, 1976; Hansen et al., 2013; Maures et al., 2014; Shi and Murphy, 2014). For example, castrated Korean eunuchs were reported to have lived 15 to 20 years longer than non-castrated men of similar socio-economic status (Min et al., 2012), while Chinese emperors known for extremely promiscuous behavior lived ∼35% shorter than their counterparts (Shi et al., 2015). *Caenorhabditis elegans* has been established as an outstanding model to investigate these interconnections. In *C. elegans*, removal of germline stem cells leads to a significant increase in lifespan (Hsin and Kenyon, 1999) as well as fat accumulation in the somatic tissues (O’Rourke et al., 2009). By contrast, mating accelerates the proliferation of germline stem cells, increases progeny production, and causes a dramatic fat loss and lifespan decrease (Shi and Murphy, 2014). The genetic analysis of mechanisms underlying the effects of mating or removal of reproductive system on *C. elegans’* fat metabolism and longevity have implicated a network that includes insulin/IGF-1 (Insulin-like Growth Factor-1) signaling, steroid signaling, lipolysis, autophagy, NHR (Nuclear Hormone Receptor) signaling, and fatty acid desaturation (Lapierre and Hansen, 2012). However, the exact relationship between fat loss and lifespan reduction is largely unknown; while fat is generally regarded as its energy source (Hansen et al., 2013), we do not know whether fat is depleted to enhance reproductive outcome, and if fat loss decreases somatic maintenance and thus shortens lifespan. It also remains unclear exactly which fatty acids are depleted in mated *C. elegans*, and whether different types of fatty acids contribute to different effects in mating-induced death.

Fatty acids are precursor molecules for all lipid classes, including storage lipids (TAGs), membrane lipids (phospholipids and sphingolipids), and signaling lipids (fatty acyl amides, eicosanoids, and others)(Fahy et al., 2009). Fat metabolism plays a crucial role in regulating the lifespan of germlineless mutants and worms with reduced reproduction. Under nutrient-poor and oxidative stress conditions, omega-3 and omega-6 fatty acids mediate the balance of lipid stores between the soma and germline (Lynn et al., 2015), suggesting a pivotal role of fatty acids in coordinating reproduction and somatic aging.

However, these previous *C. elegans* studies examined fat metabolism in animals with no or very low reproduction: either in germlineless mutants, or under conditions such as nutrient deprivation and oxidative stress where reproduction is very limited, or in self-fertilized hermaphrodites limited by sperm number. It remains unclear how fat metabolism changes when reproduction is *increased*, and whether in this circumstance fatty acids still play a role in coordinating somatic aging and reproduction. To address this question, here we increased worms’ reproduction in the most natural way by mating them with males, which typically increases progeny production by 100%-200% (Ward and Carrel, 1979; Hodgkin and Barnes, 1991), and examined lifespan, fat levels, transcriptional changes, and supplementation with fatty acids. Remarkably, oleic acid (OA) treatment specifically restores the fat loss induced by mating, and also rescues the lifespan reduction induced by mating, without affecting reproduction. This lifespan rescue by oleic acid supplementation is also pertinent to short-term mating and is conserved in gonochoristic (male and female) *C. remanei* species. Our results suggest that increased reproduction is not associated with inevitable lifespan reduction, and that metabolism of a specific fatty acid is able to uncouple the costs of reproduction from somatic longevity regulation.

## Results

### Mating induces significant lifespan decrease regardless of initial level of overall fat storage

In *C. elegans*, extended longevity is associated with altered fat metabolism and reduced reproduction, while mating significantly reduces lifespan and induces significant fat loss. However, all previous studies of the links between longevity and fat storage used worms with limited reproduction. While *daf-2* (Insulin signaling) and germlineless *glp-1* mutants are long-lived and store more fat (**Figure 1 - figure supplement 1A-C**), dietary restricted worms, such as *eat-2*, are long-lived but have less overall fat (Brooks et al., 2009; O’Rourke et al., 2009; Bar et al., 2016), making the relationship between fat levels and longevity less clear. Therefore, examining the relationship between fat metabolism and lifespan regulation in mated worms could provide new insights.

We previously showed that mating shortens lifespan and induces significant fat loss in both wild-type worms and longevity mutants, including those with excessive fat accumulation such as *daf-2* and *glp-1* (**Figure 1 - figure supplement 1A-C**, (Shi and Murphy, 2014)). We wondered whether the lifespan of longevity mutants with reduced fat storage would also be affected by mating. Dietary restriction decreases the overall fat content but robustly extends lifespan across organisms (Walker et al., 2005; Ruetenik and Barrientos, 2015; López-Otín et al., 2016). We tested the lifespan of mated worms under dietary restriction, using two methods of dietary restriction (Greer and Brunet, 2009). *eat-2* mutants have defects in pharyngeal pumping, thus consume less food than wild-type animals (Avery, 1993). Solid plate-based dietary restriction (sDR) takes the colony-forming unit of bacteria into account, using 1 × 10^11^ cfu/mL for ad libitum feeding and 1 × 10^8^ cfu/mL for dietary restriction (Greer and Brunet, 2009). We found that mating significantly decreases the lifespan of DR worms, just as it does in wild-type animals (**Figure 1A, B**). Similarly, mating also led to a significant fat loss in *eat-2* mutants (**Figure 1 - figure supplement 1D**). Thus, the reduced level of overall fat storage prior to mating in DR-treated animals does not protect the worms from mating-induced death.

**Figure 1.**
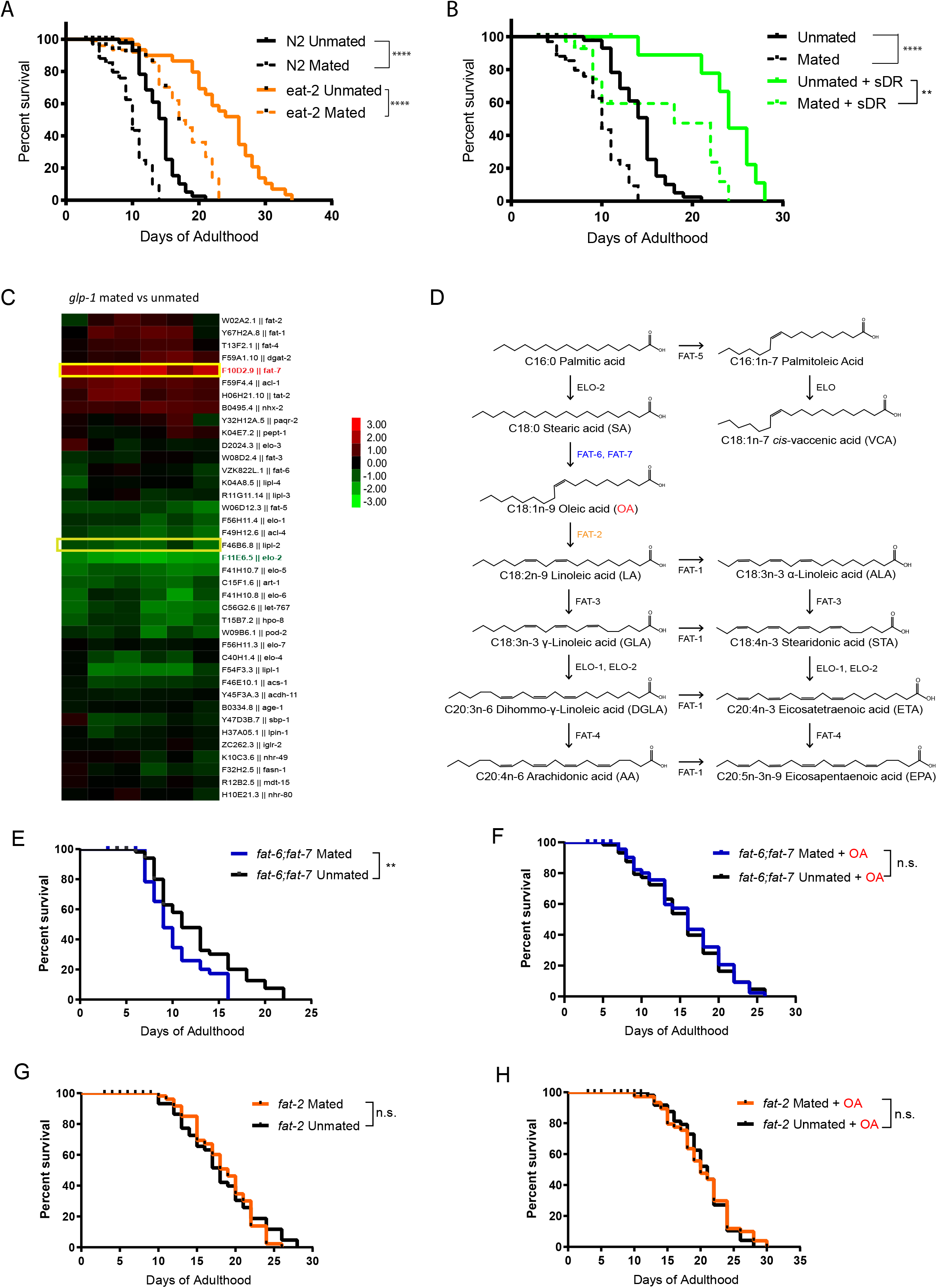
Endogenous oleic acid protects worms from mating-induced death. t test, *p <, 0.05, **p < 0.01, ***p < 0.001, ****p < 0.0001 for all graphs. For all the lifespan assays performed in this study, Kaplan-Meier analysis with log-rank (Mantel-Cox) test was used to determine statistical significance (lifespan of mated worms was always compared to unmated worms of the same genotype/treatment unless stated otherwise). Error bars represent SEM unless noted. n.s., not significant. a.u., arbitrary units. (A - B) Mated worms under dietary restriction live longer than those ad libitum despite extreme depletion of fat. N2 Unmated: 14.0 ± 0.4 days, n = 50, N2 mated: 8.8 ± 0.4 days, n = 100, p<0.0001; *eat-2(ad465)* unmated: 23.6 ± 1.1 days, n = 50, *eat-2(ad465)* mated: 15.1 ± 1.2 days, n = 100, p<0.0001; unmated + sDR: 23.7 ± 1.4 days, n = 50, mated + sDR: 14.1 ± 2.0 days, n = 100, p=0.0042.. (C) Expression heatmap of genes related to fatty acid biosynthesis in mated *C. elegans glp-1(e2141)* hermaphrodites. (D) Pathway of de novo fatty acid synthesis in *C. elegans*, including molecular structure of each fatty acid. (E,F) Mutants that cannot synthesize oleic acid have dramatic lifespan loss after mating, and oleic acid supplementation fully rescues the mating-induced lifespan decrease *fat-6;fat-7* unmated: 12.3 ± 0.7 days, n = 60; *fat-6;fat-7* mated: 9.6 ± 0.4 days, n = 100, p=0.0059; *fat-6;fat-7* unmated + oleic acid: 15.1 ± 0.8 days, n = 60; *fat-6;fat-7* mated + oleic acid: 15.5 ± 0.8 days, n = 100, p=0.7648. (G,H) Mutants that contain excess endogenous oleic acid are protected from lifespan loss after mating. *fat-2* unmated: 18.1 ± 0.8 days, n = 60; *fat-2* mated: 18.4 ± 0.6 days, n = 100, p=0.8540; *fat-2* unmated + oleic acid: 20.4 ± 0.6 days, n = 60; *fat-2* mated + oleic acid: 20.2 ± 0.7 days, n = 100, p=0.7395.

### Endogenous oleic acid protects worms against mating-induced death

To better understand how fat metabolism is altered after mating, we performed transcriptional analysis of mated vs. unmated *glp-1* hermaphrodites. *glp-1* mothers lack a germline to produce any eggs, which allowed us to disregard transcriptional changes in eggs, and instead focus on somatic changes. The lifespan of *glp-1* animals is decreased by mating and they lose over 40% of their fat stores (Shi and Murphy, 2014). Our transcriptional analysis revealed that genes with significant expressional changes in response to mating were enriched for *organic acid metabolic process* (q-value: 9.8e^-22^) and *lipid catabolic process* (q-value: 3e^-06^) (**Figure 1 - figure supplement 2**), which is consistent with the mating-induced fat loss phenotype. The lipid-regulating gene *fat-7* was significantly induced in mated worms, while *elo-2* was significantly downregulated (**Figure 1C**). Both of the genes are involved in PUFA synthesis: *fat-7* encodes a Δ9 desaturase enzyme that converts C18:0 stearic acid into C18:1n-9 oleic acid (OA), while *elo-2* encodes a key enzyme that regulates elongation of C16 and C18 fatty acids (**Figure 1D**). (There are three main types of fatty acids: saturated, monounsaturated, and polyunsaturated, which refers to fat molecules that have none, one, and more than one unsaturated carbon bond, respectively.) ELO-2 functions in several different steps of the pathway, and loss of function of *elo-2* causes multiple defects (Kniazeva et al., 2003). FAT-6 and FAT-7 are required to generate oleic acid from its direct upstream precursor stearic acid (**Figure 1D**), and act in compensatory mechanism in single mutants (Brock et al., 2006). *fat-6;fat-7* double mutants cannot endogenously synthesize oleic acid and downstream polyunsaturated fatty acids, and instead rely solely in fatty acids provided by the bacterial diet (Brooks et al., 2009); OA is almost non-existent in the common lab bacteria, OP50 (Deline et al., 2013). *fat-2* encodes the desaturase enzyme that converts oleic acid to its direct downstream product, linoleic acid (**Figure 1D**). In *fat-2* worms, oleic acid accumulates to almost a quarter of all fat, which is nearly 15-fold above the 1.7% found in wild-type animals (Watts and Browse, 2002). Therefore, these two mutants, *fat-6;fat-7* and *fat-2*, represent the effects of lack of OA and accumulation of OA, respectively.

To understand the importance of endogenous oleic acid in mating-induced death, we compared these mutants’ mated and unmated lifespans. *fat-6;fat-7* mutants displayed many visible defects, such as slow growth, low brood size, and decreased body size, and have a short lifespan relative to wild-type worms (Brock et al., 2007). Mated *fat-6;fat-7* worms lived shorter than unmated *fat-6;fat-7* worms (**Figure 1E**), indicating that mating-induced death does not require the presence of OA. Supplementation with oleic acid greatly increased the lifespans of mated *fat-6;fat-7* worms, eliminating the difference between mated and unmated animals (**Figure 1F**), suggesting that the addition of exogenous OA both increases lifespan and eliminates mating-induced lifespan reduction.

By contrast with *fat-6;fat-7* mutants, *fat-2* mutants, which accumulate oleic acid, were both long-lived prior to mating (13% increase relative to unmated N2; p=0.0045; **Figure 1 - figure supplement 3**) and resistant to mating-induced lifespan reduction (**Figure 1G**). Further supplementation of oleic acid had no effect on *fat-2* lifespan (**Figure 1H**, p=0.0915), suggesting that presence of endogenous oleic acid grants *fat-2* mutants protection from mating-induced death.

### OA specifically increases mated lifespan

Since excess endogenous oleic acid (via mutants) protects animals against mating-induced death, we wondered if supplementing mated worms with exogenous oleic acid could also mitigate mating-induced death. Therefore, we supplemented mated wild-type (N2) hermaphrodites with oleic acid or with fatty acids (FA) in the FA synthesis pathway to determine the specificity of fatty acid supplementation. Oleic acid, linoleic acid, *cis*-vaccenic acid, dihommo-y-linoleic acid, and eicosapentaenoic acid were added to the agar media of mated and unmated wild-type *C. elegans* hermaphrodites. Mated N2 worms without any fatty acid supplementation had ∼30% shorter lifespan than unmated worms (**Figure 1A-B, 2A**). While oleic acid supplementation completely rescued the lifespan loss caused by mating (**Figure 2B**), supplementation with any other fatty acid failed to rescue mating-induced lifespan decrease (**Figure 2C-F**), suggesting that oleic acid is specifically required for post-mating lifespan regulation. We supplemented the plates with various concentrations of oleic acid and found that oleic acid showed a positive lifespan effect at low concentration (0.8 mM;10% increase, p-value<0.001) and completely inhibited mating-induced death at higher concentration (2 mM; 30% increase, p<0.001) (**Figure 2G**).

**Figure 2.**
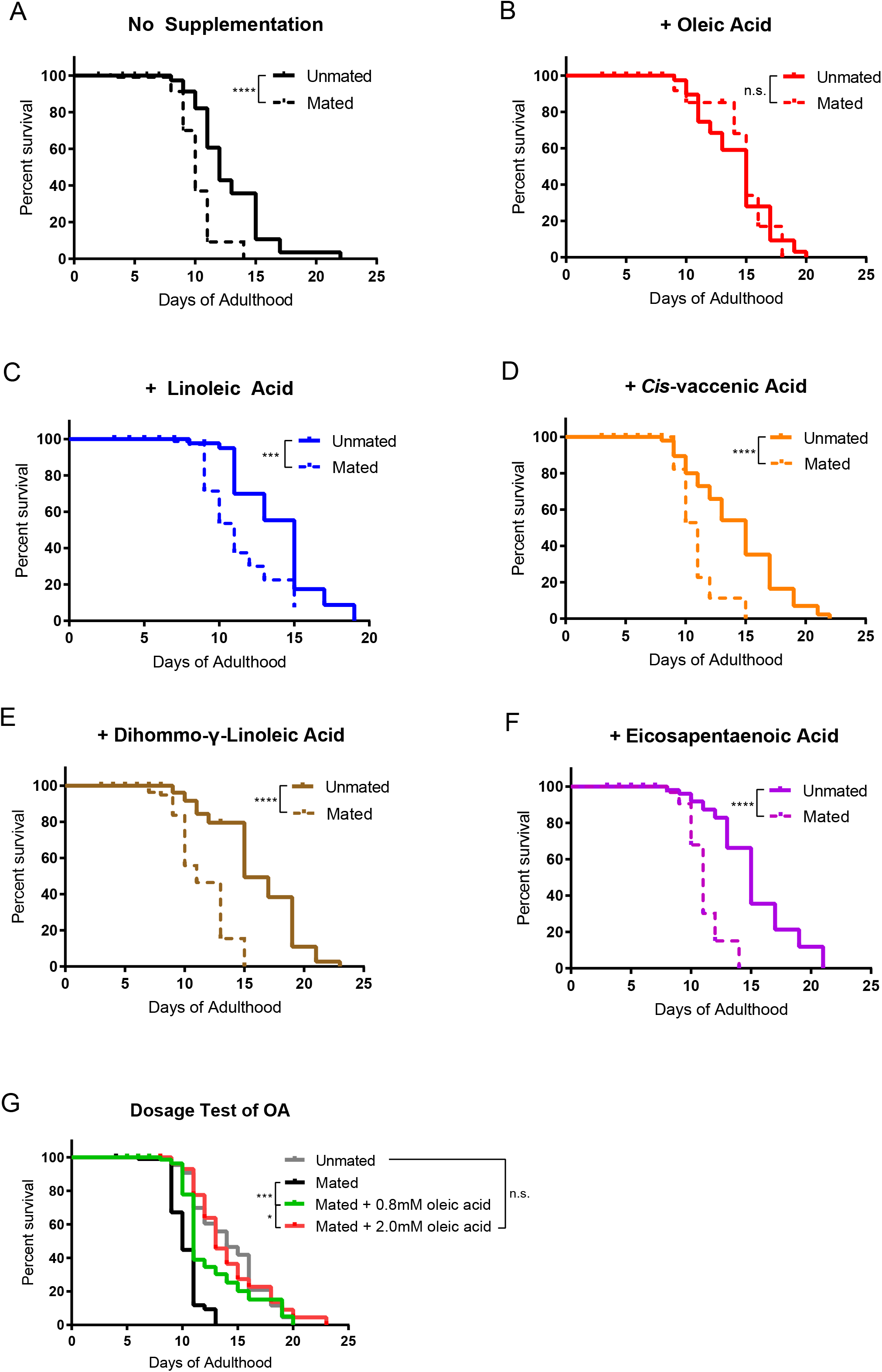
Oleic acid supplementation rescues mating-induced death in wild-type *C. elegans* mothers. (A - F) Only oleic acid supplementation rescues lifespan loss induced by mating. N2 unmated: 12.7 ± 0.6 days, n = 60; mated: 9.4 ± 0.4 days, n = 100, p<0.0001; unmated + OA: 14.2 ± 0.5 days, n = 60; mated + OA: 13.3 ± 1.2 days, n = 100, p=0.8344; unmated + LA: 14.3 ± 0.6 days, n = 60; mated + LA: 10.2 ± 0.4 days, n = 100, p=0.0007; unmated + VCA: 14.3 ± 0.6 days, n = 60; mated + VCA: 10.5 ± 0.3 days, n = 100, p<0.0001; unmated + DGLA: 15.9 ± 0.6 days, n = 60; mated + DGLA: 9.8 ± 0.5 days, n = 100, p<0.0001; unmated + EPA: 15.2 ± 0.5, n = 60; mated + EPA: 10.3 ± 0.4, n = 100, p<0.0001. (G) Dosage test of oleic acid supplementation. With improved fatty acid dissolution technique, 0.8mM of supplementation was deemed sufficient for all later experiments. Unmated N2: 14.2 ± 0.5 days, n = 50; mated: 9.5 ± 0.3 days, n = 100; mated + 0.8mM oleic acid: 12.2 ± 0.6 days, n = 100; mated + 2.0mM oleic acid: 13.8 ± 0.7 days, n = 100.

### Oleic acid supplementation restores lifespan and fat loss in mated worms without affecting reproductive health

Because lifespan, reproduction, and fat storage are closely coupled, we next checked the overall fat storage in wild-type mated and unmated worms in the presence and absence of exogenous oleic acid. We found that oleic acid supplementation fully restored mating-induced fat loss in mated worms, and also increased fat storage in unmated worms by ∼10% (**Figure 3A, B**).

**Figure 3.**
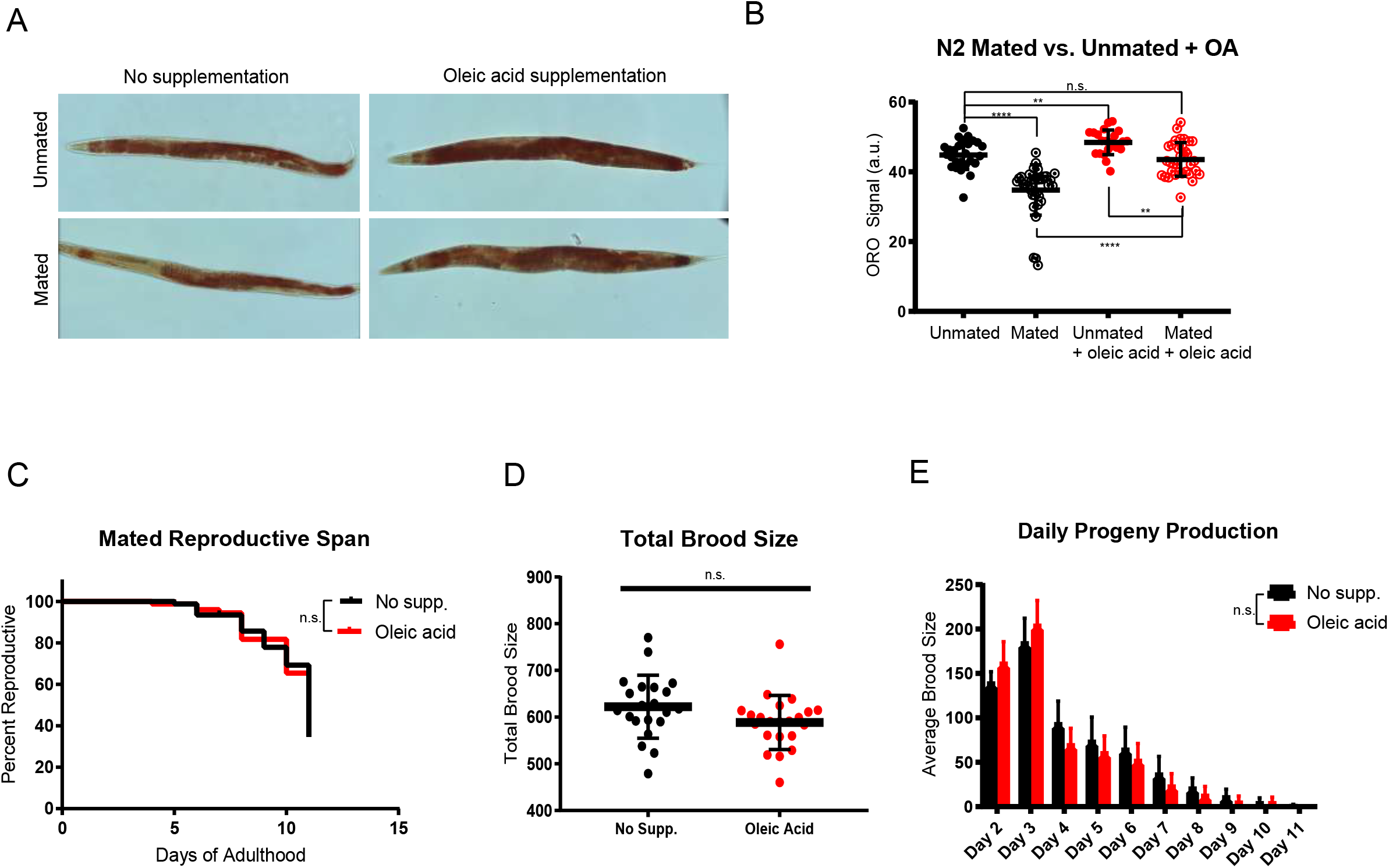
Oleic acid supplementation restores fat storage without changing the reproduction of *C. elegans* mothers. (A) Dietary supplementation of oleic acid restores overall fat storage in wild-type hermaphrodites. Representative pictures of Oil red O fat staining are shown. (B) Quantification of Oil red O fat staining. Error bars: SD. (C) Reproductive span of mated worms is not affected by oleic acid supplementation. (D) Total brood size of mated worms is not affected by oleic acid supplementation. (E, F) Daily progeny productions of mated worms with and without oleic acid supplementation are not statistically different.

Since many long-lived mutants have extended reproductive spans, we next examined whether oleic acid supplementation also extended the reproductive span of mated wild-type worms. We found that oleic acid supplementation did not have any effect on reproductive span (**Figure 3C**), total brood size (**Figure 3D**), or daily progeny production (**Figure 3E**). This result suggests that oleic acid supplementation is specifically beneficial to somatic fat metabolism and survival, without influencing reproduction.

### Lifespan rescue through oleic acid supplementation is conserved

We next wondered whether this rescue is relevant only to *C. elegans* hermaphrodites, or is conserved in non-hermaphroditic worms. We first tested *C. elegans* hermaphrodites with no self-sperm. Previously, self-sperm was found to protect and slow down mating-induced death in *C. elegans* hermaphrodites through nuclear localization of HLH-30/TFEB, a key pro-longevity regulator (Shi et al., 2019). This protective effect was confirmed in short-term mating of N2 hermaphrodites for two hours on Day 3 of adulthood, when self-sperm is still present (**Figure 4A**); two hours of mating is sufficient to induce a significant lifespan reduction in worms who lack self-sperm and therefore also lack the protective antagonist insulin (INS-37). Specifically, *fog-2* (feminization of germline) mutants lack self-sperm (Schedl and Kimble, 1988) (**Figure 4B**) and are short lived after mating. Oleic acid supplementation had no effect on lifespan on either unmated or 2hr-mated wild-type hermaphrodites that had self-sperm (**Figure 4C**), but oleic acid supplementation fully rescued the lifespan reduction of mated *fog-2* (self-spermless) mutants (**Figure 4D**). This conserved effect was further validated in *hlh-30* mutants that also suffer from 2hr-mating-induced death even in the presence of self-sperm (**Figure 4E**); oleic acid supplementation rescued *hlh-30’s* mated lifespan decrease, as well (**Figure 4F**), suggesting that oleic acid may act downstream of HLH-30/TFEB to regulate lifespan.

**Figure 4.**
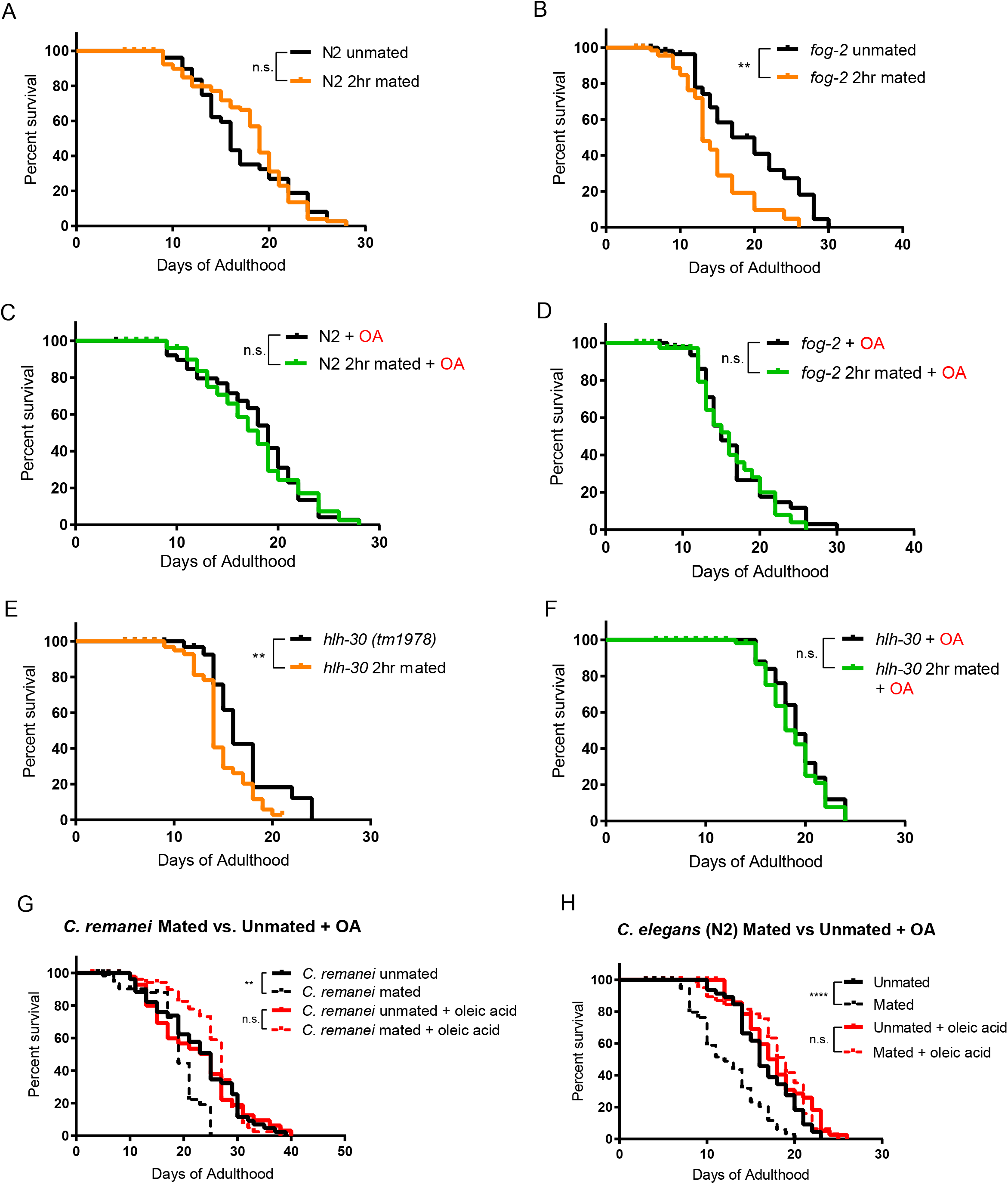
Oleic acid specific mating-induced death rescue is conserved. (A-F) Oleic acid protects worms from short-term mating-induced death. (A) unmated N2: 16.7 ± 0.8 days, n = 60; 2hr mated: 17.8 ± 0.5 days, n = 100, p=0.5900. (B) unmated *fog-2*: 18.2 ± 1.4 days, n = 60; *fog-2* 2hr mated: 14.0 ± 1.0 days, n = 100, p=0.0044. (C) unmated N2 + oleic acid: 17.4 ± 0.7 days, n = 60; 2hr mated + oleic acid: 17.7 ± 0.6 days, n = 100, p=0.7676. (D) unmated *fog-2* + oleic acid: 16.4 ± 0.8 days, n = 60; *fog-2* 2hr mated + oleic acid: 16.2 ± 0.9 days, n = 100, p=0.5910 (E) unmated *hlh-30*: 16.6 days ± 0.8 days, n = 60; *hlh-30* 2hr mated: 14.3 ± 0.4 days, n = 100, p=0.0117 (F) unmated *hlh-30* + oleic acid: 19.4 ± 0.5 days, n = 60; *hlh-30* 2hr mated + oleic acid: 18.8 ± 0.4 days, n = 100, p=0.4357. (G) Mated *C. remanei* mothers’ lifespans are also rescued by oleic acid supplementation. Unmated: 22.6 ± 1.2 days, n = 60; mated: 18.2 ± 0.8 days, n = 60, p=0.0029; unmated + oleic acid: 21.9 ± 1.4 days, n = 60; mated + oleic acid: 25.3 ± 0.9 days, n = 60, p=0.3266. (H) Lifespans of unmated and mated N2 hermaphrodites with and without oleic acid supplementation. Unmated: 16.6 ± 0.5 days, n = 49; mated: 11.8 ± 0.5 days, n = 80, p<0.0001; unmated + oleic acid: 17.9 ± 0.6 days, n = 50; mated + oleic acid: 18.1 ± 0.6 days, n = 80, p=0.9466.

Finally, we wondered if oleic acid supplementation is specific for hermaphrodites, or whether it can also rescue the early death of mated gonochoristic species females. We showed previously that *C. remanei* females also have significantly reduced lifespan after mating with *C. remanei* males (Shi and Murphy, 2014). Remarkably, mated females supplemented with oleic acid lived as long as unmated females with and without supplementation (**Figure 4G**), just as in the case of *C. elegans* wild-type hermaphrodites (**Figure 4H**). These results suggest that lifespan rescue through oleic acid supplementation is evolutionarily conserved in mated *Caenorhabditis* mothers.

## Discussion

The trade-off between reproduction and lifespan has been observed in a variety of animals (Partridge et al., 2005; Hansen et al., 2013). Fat, as the major means of storing energy, has been shown to be closely involved in both reproduction and lifespan regulation. The same inverse relationship also exists between reproduction and fat storage (Hansen et al., 2013). Germline-less *glp-1* worms upregulate the Δ9 desaturase enzyme FAT-6/SCD1 (Stearoyl-CoA Desaturase 1)(Goudeau et al., 2011) and shift the lipid profile to increase somatic maintenance and longevity (Ratnappan et al., 2014), and deficiency of the H3K4me3 methyltransferase (*ash-2, set-2*) upregulate Δ9 desaturases FAT-5 and FAT-7, leading to the accumulation of mono-unsaturated fatty acids and subsequent lifespan extension (Han et al., 2017).

By contrast, mated hermaphrodites demonstrated the inverse relationship between reproduction, longevity, and fat storage: mating increases progeny production by 2-3 fold, reduces fat stores, and significantly shortens lifespan compared to unmated hermaphrodites. Here, we have shown that supplementing the worms with oleic acid can completely rescue the shortened lifespan and reduced fat store of mated mothers, whereas supplementing other fatty acids is not able to ameliorate the mating-induced death. Dietary supplementation of oleic acid as well as a few other MUFAs including palmitoleic and cis-vaccenic acid can extend the lifespan of unmated hermaphrodites (Han et al., 2017). Taken together, our results demonstrate the specificity of oleic acid in regulating post-mating lifespan, and suggest that it is slightly different from H3K4me3 methyltransferase-mediated transgenerational inheritance of longevity.

Our analysis of mated animals with increased reproduction revealed that overall (total) fat levels plays no role in the lifespan regulation of mated worms; that is, mating reduces the lifespan of longevity mutants, whether they exhibit increased or decreased fat levels prior to mating. Our transcriptome analysis of mated and unmated worms revealed mating-induced upregulation of regulators of fatty acid metabolism. Only oleic acid treatment can rescue fat loss induced by mating and lifespan reduction induced by mating, without affecting reproduction. This lifespan rescue by oleic acid supplementation is also pertinent to short-term mating and is conserved in gonochoristic (male and female) *C. remanei* species. Here, we have shown that oleic acid, a product of the Δ9 desaturase enzyme FAT-7 - whether through its endogenous accumulation by loss of FAT-2 activity or through exogenous supplementation - can prevent mating-induced death (**Figure 5**). Furthermore, the lifespan-rescuing effect of oleic acid is evolutionarily conserved in self-spermless hermaphrodites and females of gonochoristic *Caenorhabditis* species. Our results suggest that oleic acid depletion in the soma is the ultimate downstream cost of reproduction. Oleic acid might affect the soma through maintenance of membrane fluidity (stearic acid/oleic acid ratio) or because of general lipid storage. It is notable that other lipids in the pathway were not able to rescue mating-induced death, suggesting that this specific fatty acid has a unique role in somatic maintenance and growth.

**Figure 5.**
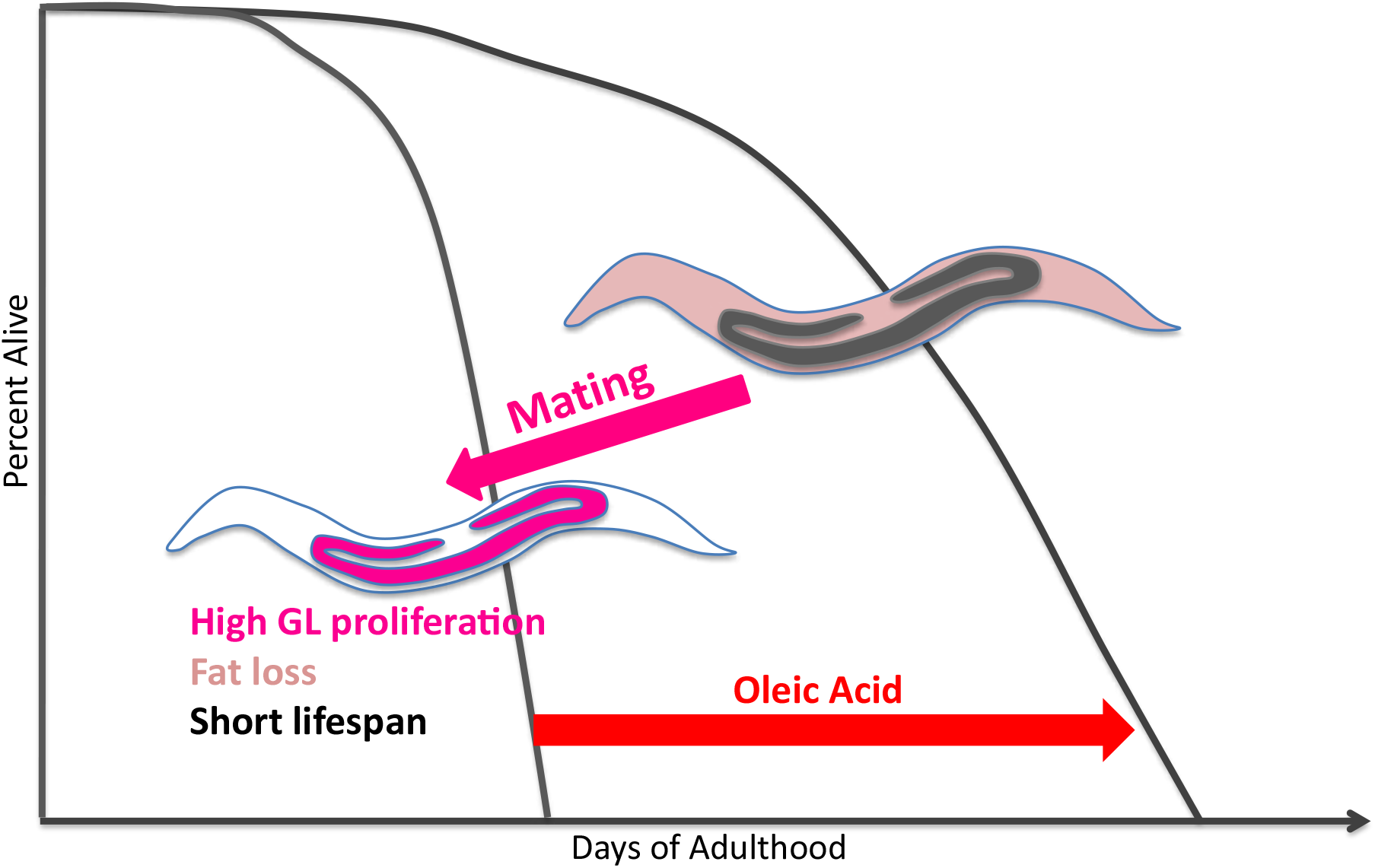
Model of oleic acid-mediated lifespan restoration in mated worms. Mating leads to germline hyperactivity, fat loss, and shortened lifespan in *Caenorhabditis* mothers. Oleic acid additon (whether endogenously or exogenously) is able to restore the lifespan decrease caused by mating.

The specificity of oleic acid has also been reported in mediating the age-dependent somatic depletion of fat (Asdf) (Lynn et al., 2015), as oleic acid is the only fatty acid that is able to rescue the Asdf phenotype. Even though the Asdf phenotype was examined in unmated worms and under oxidative stress, which differs from the mating-induced death in this study, both phenotypes involve the interaction between the soma and germline. The fact that the specificity of oleic acid is observed in both studies further increases the likelihood of oleic acid as the possible link between germline/reproduction and somatic maintenance/lifespan.

In addition to dietary supplementation, we also show that endogenous oleic acid is beneficial to mated worms. Genetically blocking FAT-2, the desaturase enzyme that converts oleic acid to linoleic acid, increases endogenous oleic acid levels (Watts and Browse, 2002), which we find protects the worms from mating-induced death. FAT-6 and FAT-7 are the stearoyl CoA desaturases (SCD) that convert stearic acid to oleic acid. Interestingly, they are up-regulated in long-lived longevity mutants such as *daf-2* and *glp-1* (Murphy et al., 2003; Goudeau et al., 2011), indicating again that increased endogenous oleic acid is beneficial to worms. It is possible that oleic acid is the somatic energy source that extends lifespan in these mutants. FAT-6/FAT-7/SCD1 desaturase enzyme activity is highly conserved. In mammals including humans, oleic acid is the prime substrate for triglyceride (Chu et al., 2006). Loss of SCD1, which synthesizes oleic acid, was shown to protect against adiposity by preventing accumulation of triglyceride in mice (Ntambi et al., 2002), just as *fat-6;fat-7* mutant worms have significantly low levels of triglyceride storage (Watts and Browse, 2002). Furthermore, SCD1 is strongly upregulated in response to ovariectomy in mice (Paquette et al., 2008). The biological roles of oleic acid and desaturation are thus conserved in several aspects between *C. elegans* and humans. With regard to reproduction, oleic acid is a major determinant of plasma membrane fluidity in many cells, including follicular cells (Funari et al., 2003). In bovine oocytes, oleic acid plays a key role in membrane fluidity that counteracts the detrimental effects of exposure to saturated fatty acids (Aardema et al., 2011), and acts as metabolic regulator of oxidative stress and cellular signaling in oocytes, early embryonic development, and female fertility (Fayezi et al., 2018).

Oleic acid is likely to be the linchpin of the reproductive and somatic energy balance. Mating-induced up-regulation of *fat-7*, which converts stearic acid to oleic acid, could be a response of the mated worms to prepare themselves with more endogenous oleic acid for dramatically increased progeny production. At the same time, the somatic reserve of oleic acid is depleted as indicated by the loss of fat storage to fuel the mating-induced elevated reproductive activity. In this case, mated worms prioritize oleic acid for reproduction, and the outcome is increased reproduction at the cost of lifespan. By contrast, in *daf-2* and *glp-1* mutants with reduced or no reproductive activity, more oleic acid accumulates in the somatic tissues and may be utilized for somatic maintenance, leading to an extension of lifespan. Therefore, oleic acid is likely to be the limiting energy source that reproduction and somatic maintenance compete for. Our results suggest that oleic acid depletion is a prime example of the cost of reproduction. Excitingly, it is one that appears to be reversible through dietary supplementation of oleic acid. Oleic acid, along with the evolutionarily conserved enzymes in the pathway, might be therapeutic targets that could help humans improve reproductive and somatic health at the same time without having to sacrifice one for the other.

## FIGURE LEGENDS

**Figure 1 - figure supplement 1.**
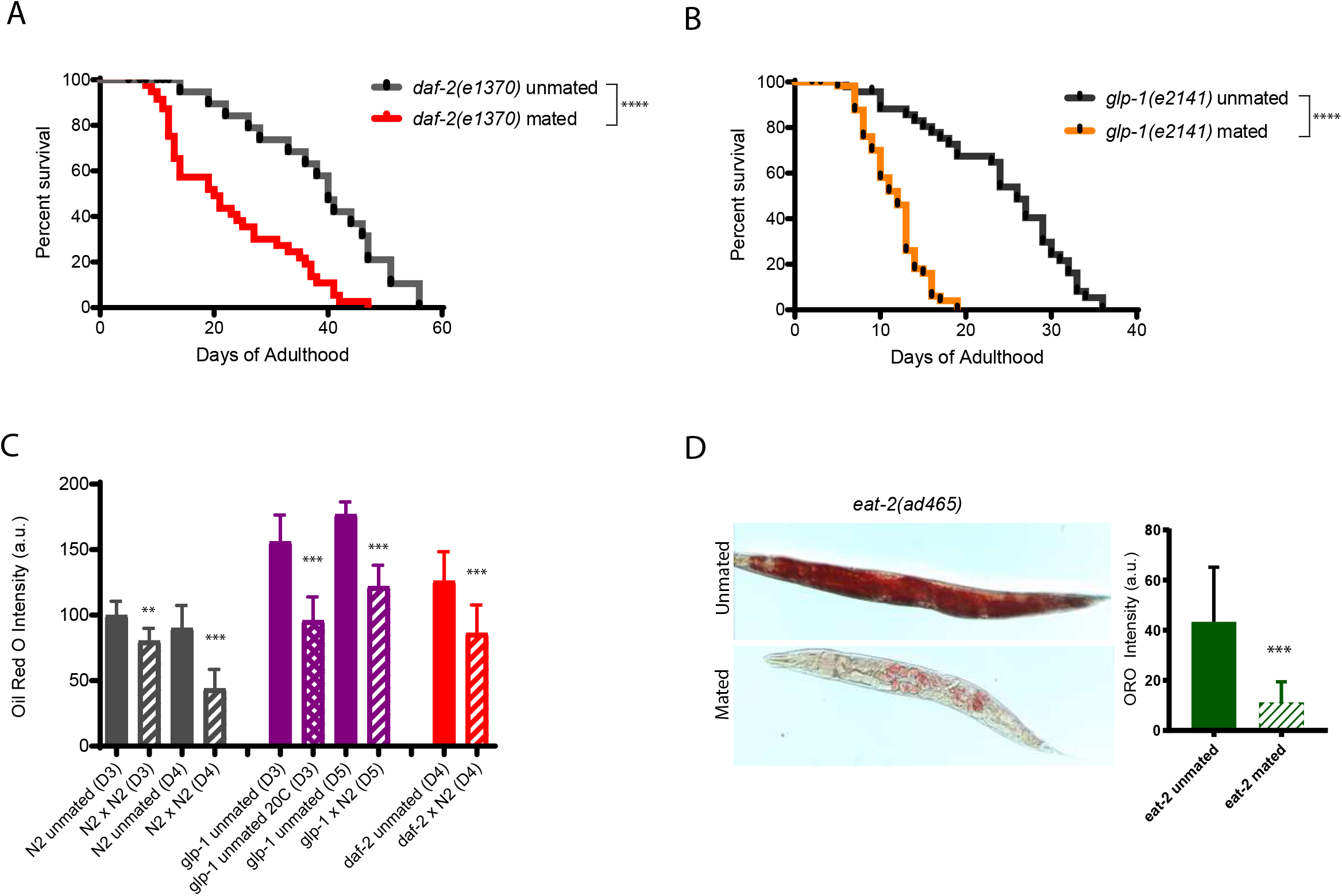
Mating causes dramatic fat loss and lifespan decrease in longevity mutants. (A) *daf-2(e1370)* longevity mutants are short-lived post-mating. Unmated *daf-2*: 37.0 ± 1.9 days, n=47, mated *daf-2*: 19.4 ± 2.2 days, n=60, p<0.0001. (B) *glp-1(e2141)* mutants live shorter after mating as well. Unmated *glp-1*: 24.1 ± 1.3 days, n=48, mated *glp-1*: 11.6 ± 0.5 days, n=60, p<0.0001. (C) Quantification of Oil red O fat staining of mated and unmated wild-type(N2), *daf-2*, and *glp-1* worms. (D) Mating also induces significant fat loss in *eat-2(ad465)* hermaphrodites. Left: representative Oil Red O staining pictures; Right: quantification of Oil Red O staining.

**Figure 1 - figure supplement 2.**
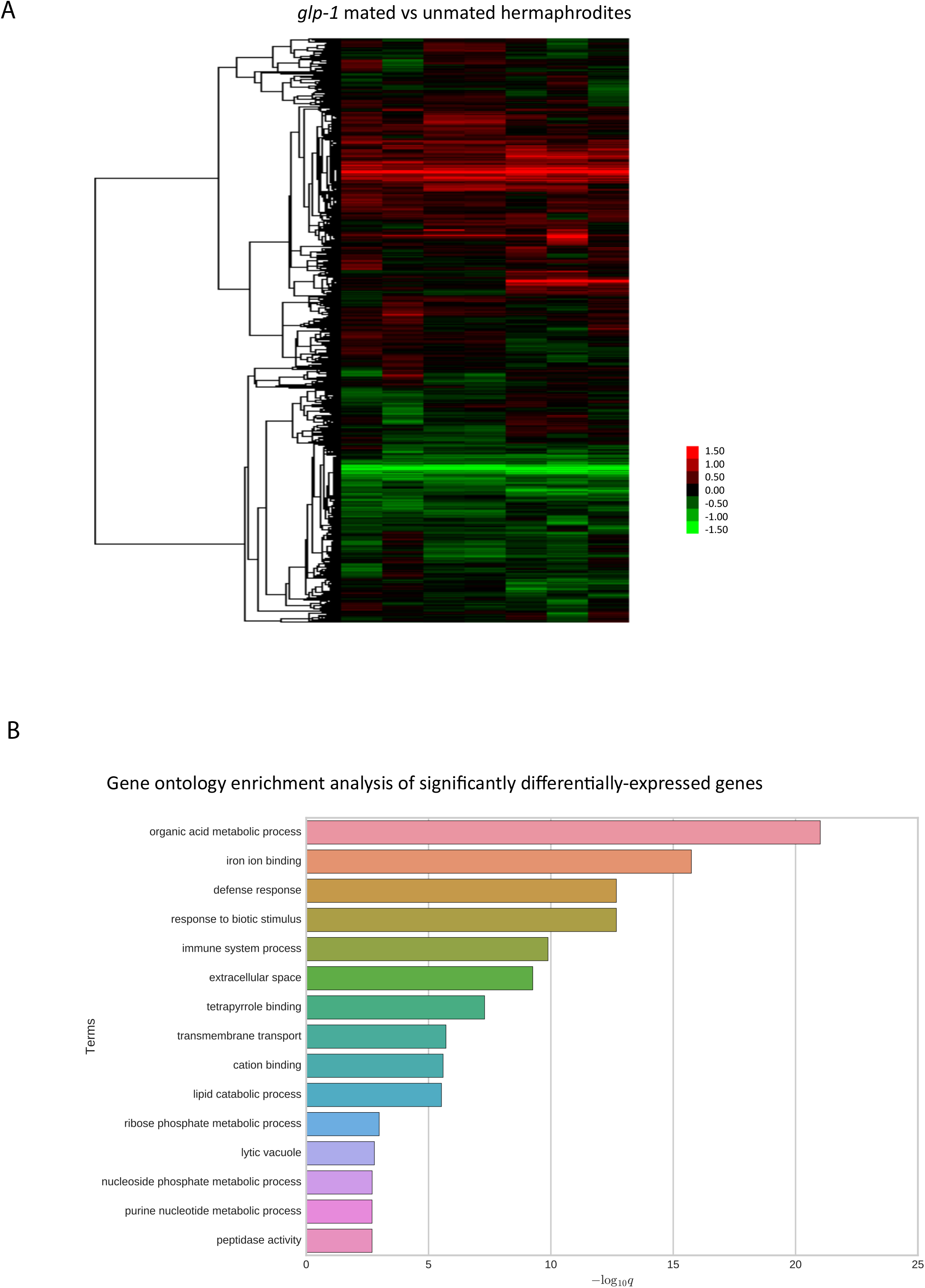
Genome-wide transcriptional analysis of mated vs unmated *glp-1* hermaphrodites. (A) Heatmap of whole transcriptome comparison between mated and unmated *glp-1(e2141)* hermaphrodites. (B) Gene ontology enrichment analysis of significantly differentially-expressed genes.

**Figure 1 - figure supplement 3.**
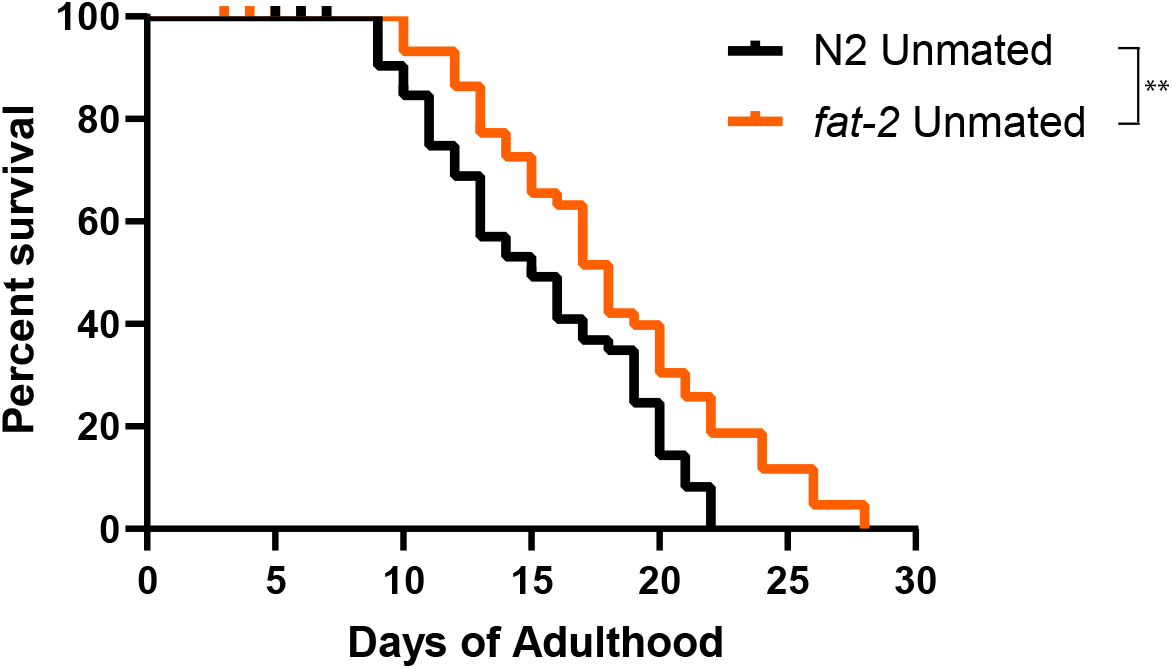
Worms with excessive endogenous oleic acid are longer-lived. Lifespans: N2 unmated: 15.4 ± 0.6 days, n = 60; *fat-2* unmated: 18.2 ± 0.8 days, n = 60, p=0.0045.

## Materials and Method

### Strains and maintenance

All worm strains used in this study are listed below. All strains were maintained on Nematode Growth Media (NGM) plates at 20°C according to the standard procedure (Brenner, 1974).

| Strain | Genotype | Source |
| --- | --- | --- |
| N2 | Wild-type | CGC |
| CB4108 | *fog-2(q71) V* | CGC |
| CB4037 | *glp-1(e2141) III* | CGC |
| RB969 | *fat-2(ok873) IV* | CGC |
| BX156 | *fat-6(tm331) IV;fat-7(wa36)V* | CGC |
| JIN1375 | *hlh-30(tm1978) IV* | CGC |
| PB4641 | *Caenorhabditis remanei* | CGC |

CGC is the *Caenorhabditis* Genetics Center

### Mating assays

All assays were performed at 20°C, and all worms were synchronized through bleaching adult worms with a mixture of NaOCl and KOH, leaving only the eggs.

1) 24hour group mating

10cm NGM plate was seeded with 300µL of OP50 (Brenner, 1974) to create a ∼3cm diameter of bacterial lawn. 100 hermaphrodites and 300 males at late L4 were transferred onto the plate. The ratio of 1 hermaphrodite to 3 males were used to optimize mating. 24 hours later, the hermaphrodites were transferred onto newly seeded plates without males. Complete mating means that males transfer enough sperm so that all progeny produced are cross-progeny (half males, half hermaphrodites). Incomplete mating means that males do not transfer enough sperm so that their sperm is used up and hermaphrodites later revert to using their own sperm, evidently seen from the ratio of male: hermaphrodite offspring. The progeny of the mated hermaphrodites was checked every day, and incompletely mated worms were censored. For *C. remanei*, same method was used for mating. Since males deposit copulatory plugs onto females’ vulva after mating, mated females without the plugs and unmated females with the plugs were both censored.

For *C. elegans* males, males and hermaphrodites were kept together for 2 days for mated condition.

2) 2hr short-term mating

Same setup was used as group mating, but the mating was allowed for only 2 hours. Hermaphrodites were transferred first onto seeded plates, and males were moved onto the plates later in an orderly and timely manner, so that each experimental group has exactly 2 hours for mating.

### Lifespan assay

Lifespan assays were performed using 3cm NGM plates seeded with 25µL of OP50. Maximum of 20 worms were present at each plate. For mated group, n = 100 while for unmated group, n >50. Worms were transferred every day to freshly seeded plates during reproductive period to eliminate confounding progeny, and every other day afterwards. Worms were scored dead if they did not respond to repeated prods with a pick. Beginning of Day 1 of adulthood was defined as t = 0, and Kaplan-Meier survival analysis was performed with the log-rank (Mantel-Cox) method. Lifespan curves were generated using GraphPad Prism. Bagged (matricide), exploded, or lost worms were censored.

*fat-6;fat-7* mutants had suspended growth, so they were considered Day 1 adults when the egg-laying began.

### Reproductive span and brood size assay

Individual hermaphrodites after mating were transferred onto 3cm NGM plates seeded with 25µL of OP50 and moved to fresh plates daily until reproduction ceased for at least two days. The old plates were left at 20°C for two days to allow the offspring to grow into adults, which were manually counted for daily production and total brood size. At least 20 plates of individual worms were counted to account for individual variation. The last day of reproduction was marked as the last day of progeny production, evidenced by no worms on the old plates. When matricide occurred, the worm was censored from the experiment on that day. Kaplan-Meier analysis with log-rank (Mantel-Cox) method was performed to compare the reproductive spans. Reproductive span curves and brood size values were graphed using GraphPad Prism.

### Fatty acid supplementation

The fatty acid supplementation was adapted from the published protocol(Deline et al., 2013) with minor modifications from a more recent publication(Han et al., 2017). The detergent Tergitol (type NP-40, Sigma-Aldrich) was added to a final concentration of 0.001% in liquid NGM media, and optimized concentration of 0.8mM was supplemented for all fatty acids. For our purpose, fatty acids were in ethanol solution and were further dissolved in purified water by shaking for 30 minutes. Working stock was made fresh every time. The media was stirred for at least 5 minutes after the working stock was added. The fatty acids were deemed incompletely dissolved if similar number of oil bubbles can be seen in the media with and without the detergent. Plates were kept inside a dark plastic box with no ventilation, but with sterile lab tissues that were replaced every day to absorb moisture. This method enabled the plates to remain without visible blackening of the fatty acid bubbles for longer period. Control plates were made using the exact same procedure including the detergent and the ethanol, but not the fatty acid. Correct and consistent application of this procedure was essential for replicability of the results.

### Oil Red O staining and quantification

Oil Red O staining was adapted from the published protocol to stain around 100 worms per condition(Wählby et al., 2014). Approximately 10 worms were imaged per slide with Nikon Ti. Oil Red O quantification was performed as published(O’Rourke et al., 2009). Briefly, colored images were split into RGB monochromatic images in Image J. The polygon selection tool was used to outline the boundary of each worm. The Oil Red O staining arbitrary unit (a.u.) was determined by mean grey value within the worm, calculated by the difference between the signals in Blue/Green and Red channel. T-test analysis was performed to compare the fat staining of different groups of worms. Individual intensity values were graphed using GraphPad Prism.

### Microarrays

*glp-1(e2141)* hermaphrodites grown at the restrictive temperature (25°C) were mated on day 1 of adulthood for 24 hr in a 2:1 male:hermaphrodite ratio. 200 hermaphrodites were collected on day 2/3 of adulthood for each biological replicate. RNA was extracted by the heat vortexing method. Two-color Agilent microarrays were used for expression analysis. Significantly differentially-expressed gene sets were identified using SAM (Tusher et al., 2001). Gene ontology enrichment analysis was performed using wormbase enrichment tools (https://wormbase.org/tools/enrichment/tea/tea.cgi).

Original microarray dataset can be found in PUMAdb: https://puma.princeton.edu/cgi-bin/exptsets/review.pl?exptset_no=7345

### Lifespan assays summary

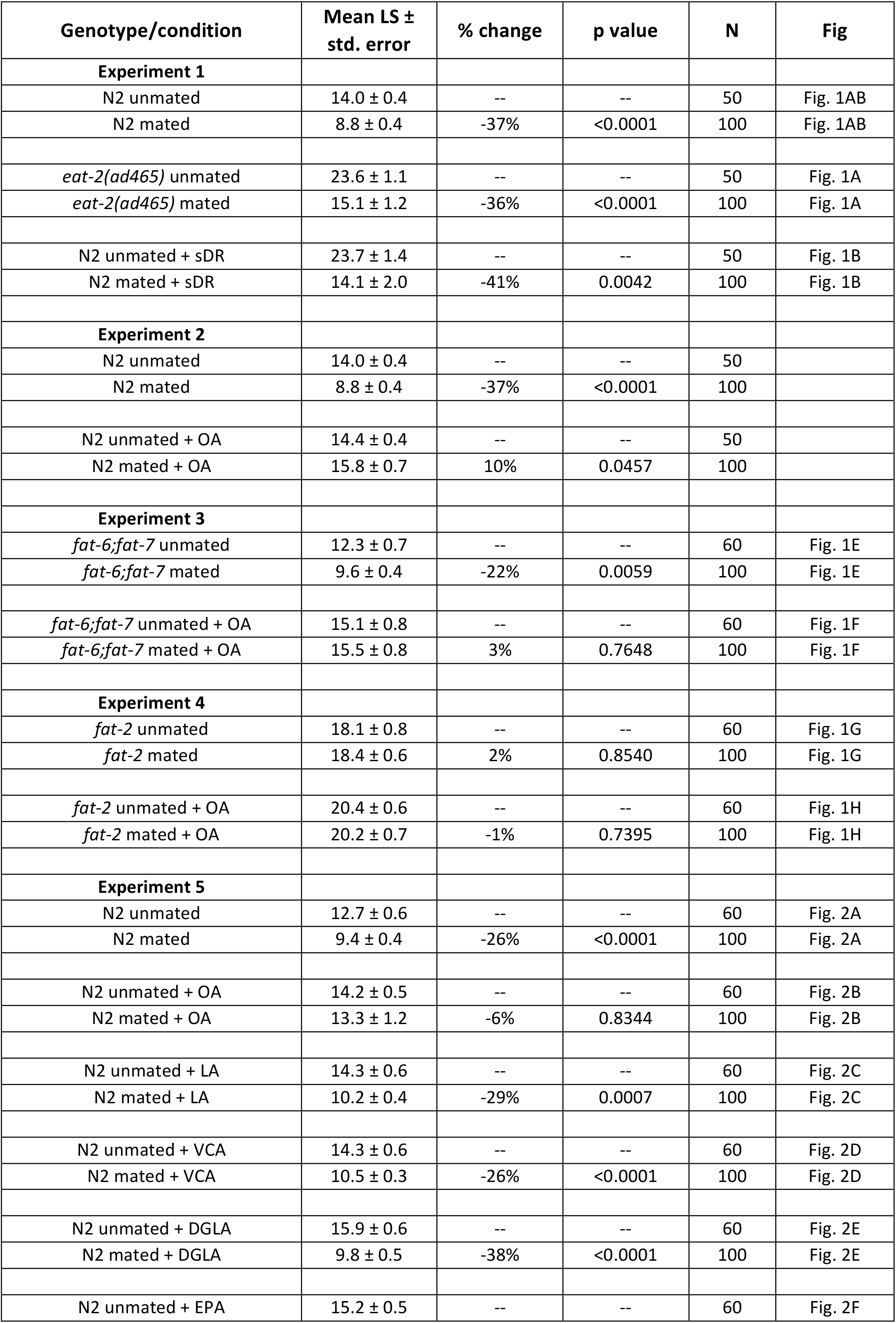
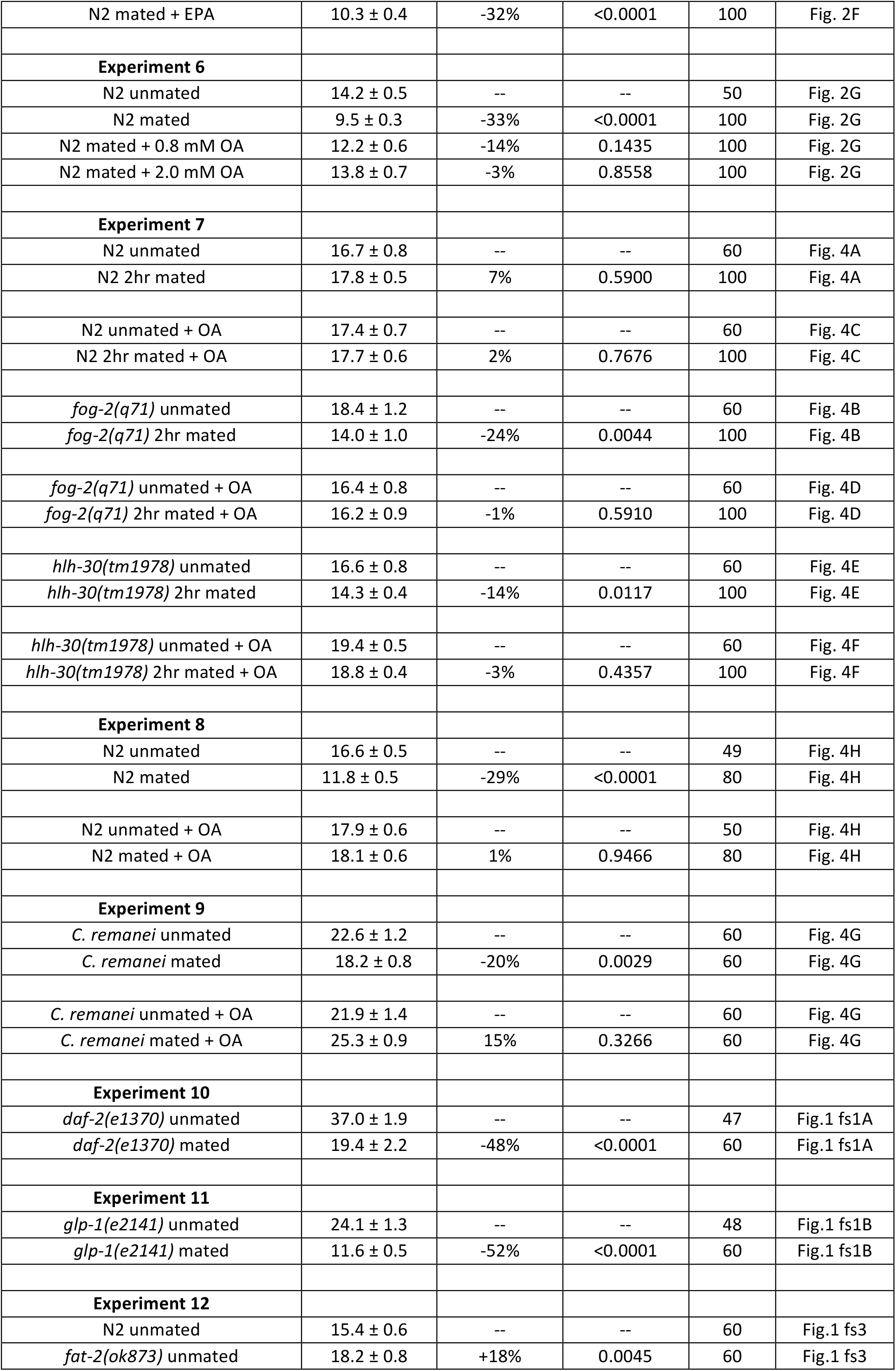

